# The Neural Basis of Attentional Blink as a Selective Control Mechanism in Conscious Perception

**DOI:** 10.1101/2024.03.30.587354

**Authors:** Q. Xin, S. Forman, K. L. Christison-Lagay, C. Micek, S. I. Kronemer, M. Aksen, L. Grobois, V. Contreras Ramirez, A. Khalaf, D. Jin, S. Aerts, M. M. Chun, M. J. Crowley, H. Blumenfeld

**Affiliations:** Interdepartmental Neuroscience Program, Yale University; New Haven, CT, USA; Department of Neurology, Yale School of Medicine; New Haven, CT, USA; Department of Neuroscience, Yale School of Medicine; New Haven, CT, USA; Department of Neurosurgery, Yale School of Medicine; New Haven CT, USA; Child Study Center, Yale School of Medicine; New Haven, CT, USA; Department of Psychology, Yale University; New Haven CT, USA

## Abstract

Conscious perception of visual stimuli involves large-scale brain networks with multiple activation-deactivation dynamics. Previous works have shown that early detection networks may be switched off about 200ms to 300ms after presentation of a visual stimulus. We hypothesize that these deactivations represent a selective control mechanism of the brain to conserve resources for post-perceptual processing. To this end, we used attentional blink as a behavioral measure for this mechanism. We showed that attentional blink is more likely to occur when a previous visual stimulus was consciously perceived. Using high-resolution eye-tracking, we found prolonged decrease in pupil diameter and transient decrease in blink probability associated with attentional blink. Using scalp EEG data, we further showed that attentional blink is associated with more pronounced event-related potentials related to visual processing and report.

**One sentence summary:** attentional blink may represent a selective control mechanism of neural processing resources underlying conscious perception.

## Main Text

Conscious perception of transient sensory stimuli has broad scientific and clinical implications. In particular, being consciously aware of transient visual stimuli is necessary for normal quality of life. Meanwhile, an impairment of such processes is also implicated in attentional disorders and epilepsy[1] and can be considerably debilitating. Previous works have shown that conscious perception elicits broad networks through a series of activation and deactivation in the first 1000ms following a sensory stimulus[2, 3]. We have proposed a model of conscious perception, the “detect”, “pulse”, “switch”, and “wave” (DPSW) framework. This framework is detailed in Blumenfeld, 2023[4]. Briefly, following a sensory stimulus, the brain first recruits the detection networks, such as primary visual cortex. Then, a pulse of signals from the subcortical arousal networks is then sent to the cortex to trigger downstream processes. This is followed by the switching off the competing processes, such as the detection network or the default mode network. Finally, a wave of cortical processing through the cortical hierarchy is present, likely for memory and report.

In this work, we focused on the “switch” aspect of the processing for two reasons. First, while evidence for this process were found in patients undergoing intracranial EEG[2], it has not been found in neurotypical subjects. Second, there lacks a behavioral outcome that can be related to the deactivation of brain networks. To address this gap in knowledge, we integrated a classical phenomenon, attentional blink, into the conscious perception paradigm. Attentional blink generally refers to the inability to perceive the second stimulus if a pair of visual stimuli are presented within 300ms from each other, as if the attention “blinks”. Specifically, we posit that the switch process reflects a fundamental mechanism of selective control of cortical activities to ensure the proper cascade of neuronal processing during conscious perception. We hypothesize that (1) conscious perception of a visual stimulus will impede the processing of a second stimulus if the second stimulus is presented within the “switch” window; and (2) conscious perception elicits similar neuronal processes as attentional blink.

To this end, we developed a paradigm to assess *perception* for the first stimulus (“T1”) and *detection* for the second stimulus (“T2”, Fig. 1A). For T1, the target was a face image with its opacity titrated to 50% of each subject’s individual perceptual threshold (see **Methods**). Each subject was asked to report (1) if they have seen the face and (2) if they can correctly identify the location of the face (1 out of 4 possible locations). To control for false positives, 12.5% of the T1 images are blanks. The time difference between T1 and T2 is referred to as “stimulus onset asynchrony” (SOA). Our tested SOA include 100ms, 200ms, 300ms, and 600ms. For T2, the target was the letter “X” embedded in a rapid serial visual presentation of other distractor letters (Fig. 1A). Each subject was only asked to report if they have detected the T2. In 20% of the trials, the target T2 letter was never shown in order to control for false positives. Concurrent scalp EEG (1000Hz, 256-channel, EGI) and eye-tracking (1000Hz, SR Research) data were collected in a subset of these subjects (n=36).

**Figure 1.**
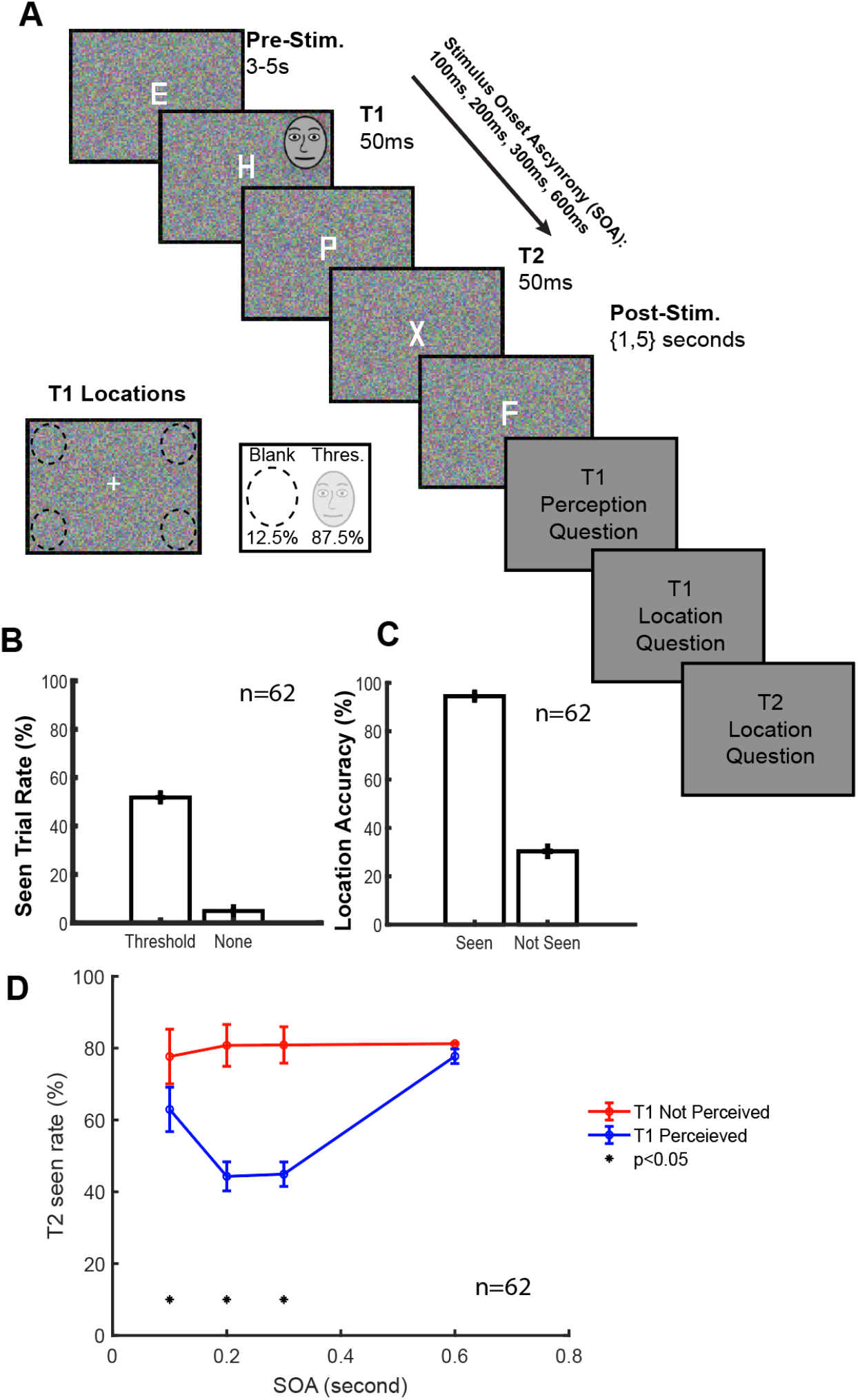
Behavior performance on T1 and T2. (A) Illustration of an example trial sequence. The participants were instructed to fixate at the center of the screen, where a stream of rapid serial visual presentation (RSVP) of letters appear for 50ms each. T1 target is a face, with opacity titrated to their 50% perceptual threshold. T2 target is the letter “X”. (B) Subject performance on the rate of T1 reported being seen for threshold and blank T1 stimuli. (C) Subject performance on the rate of correctly identifying the location of T1 when T1 reported being seen versus not seen. (D) Modulation of the rate of T2 reported being seen for different stimulus onset asynchrony (SOA) conditions. *denotes two-sample t-test <0.05 between T1-percieved and T1-not-perceived conditions. The face stimulus (**A**) is sourced from the FACES database[6].

We found that the conscious perception of T1 selectively impeded the detection of T2, thus reproducing the attentional blink effect during the switch window. Specifically, we first found that introducing T2 did not affect the performance of T1. Subjects reported a seen rate of 51.79% (±0.91%, n=62) on T1 when T1 was presented, consistent with the expected 50% perceptual threshold (Fig. 1B). In turn, when no T1 was presented (blanks), the subjects reported a seen rate of 4.90% (±0.60%, n=62), suggesting a low level of false positives (Fig. 1B). Furthermore, when subjects reported they saw T1, their accuracy in reporting the correct location was 94.49% (±0.65%, n=62; Fig. 1C). In turn, when they reported they did not see the face, their location accuracy was 30.31% (±1.1%; n=62; Fig. 1C). These results are consistent with previous studies on T1 performance, neurotypical adults and patients with neurostimulator implants (Supp. Fig. 1). Notably, the detection of T2 was significantly modulated by T1. When T1 was not perceived, regardless of SOA, the subjects reported having seen T2 at around 80% across the 4 SOA conditions, consistent with the percentage of T2 blanks (Fig. 1D). However, the T2 seen rate was significantly lower when T1 was perceived at 100ms, 200ms, and 300ms SOA conditions compared to when T1 was not perceived (100ms: p=0.0189; 200ms: p<0.0001; 300ms: p<0.0001). This difference was significant in each of these conditions. In turn, when SOA is at 600ms and outside the range of attentional blink/switch, this modulation effect was no longer present (Fig. 1D).

We found that pupil diameter and eye-blink probability were significantly modulated by attentional blink. Given T1 is perceived, pupils were significantly less dilated when attentional blink occurred compared to when no attentional blink occurred (Fig. 2A). This statistically significant difference emerged after about 1000ms since T1 onset and maintained throughout the period of 4 seconds after T1 onset. In turn, we observed a much more transient different in blink probability. Given T1 is perceived, subjects were much less likely to blink when attentional blink compared to when attentional blink did not occur (Fig. 2B). In contrast to pupil dilation, this statistically significant difference emerged earlier at about 750ms since T1 onset and only persisted for about 500ms. Afterwards, the blink probability dynamics remain the same regardless of whether attentional blink occurred.

**Figure 2.**
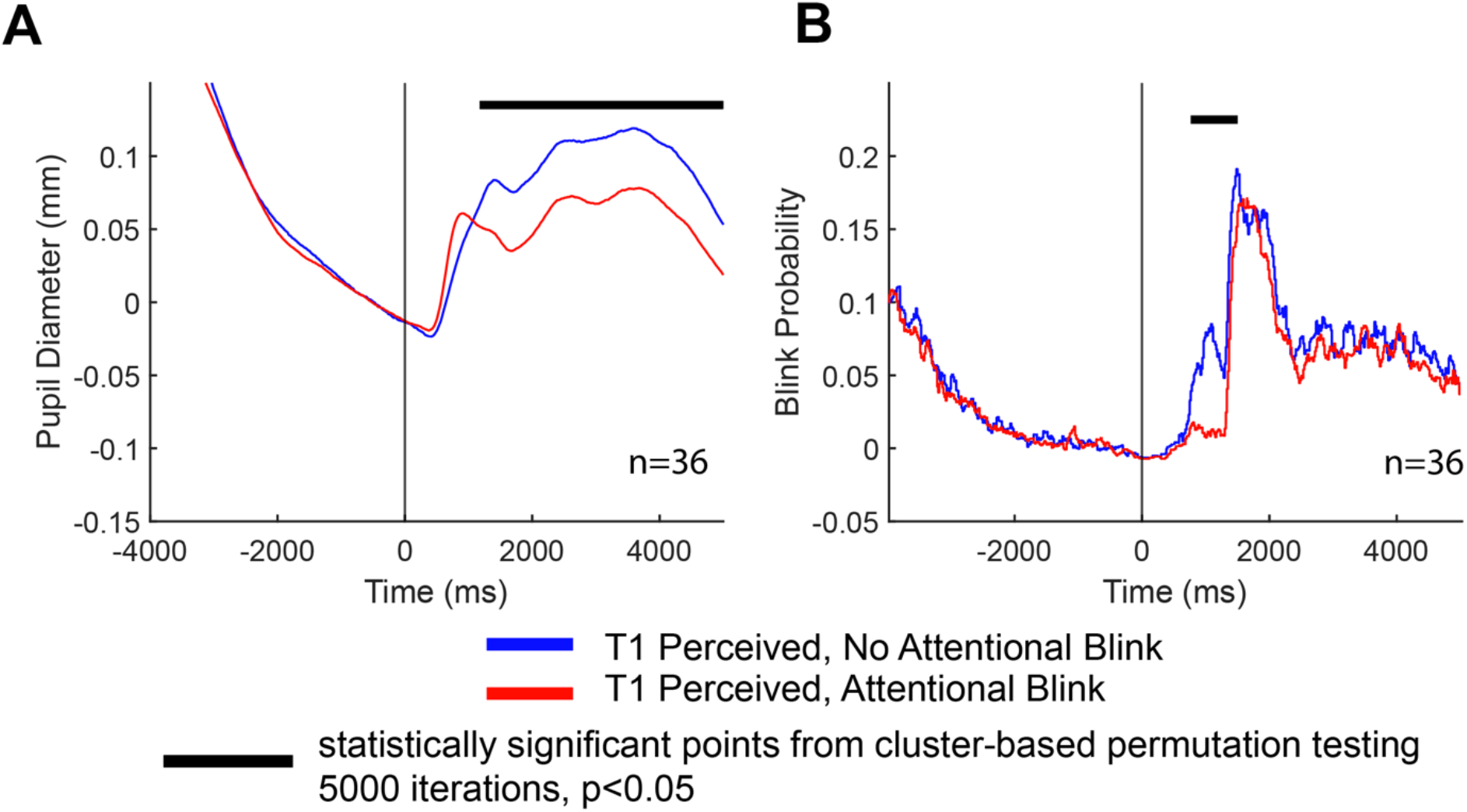
Eye-movement dynamics with respect to attentional blink. Comparison of diameter (A) and blink probability (B) between whether attentional blink occurs, given that T1 is perceived. Black bar indicates statistically significant points from cluster-based permutation testing against baseline (5000 iterations, p<0.05).

We also found that attentional blink selectively introduced differential event-related potentials in the scalp EEG responses. As illustrated by an example electrode pz (Fig. 3A), we found attentional blink led to significantly more pronounced N100, VAN, and most consistently, P300. Examining the voltage-space topoplots, it can be seen that, compared to when no attentional blink occurs (Fig. 3B), attentional blink led to a more pronounced N100, VAN, and P300 (Fig. 3C). Subtracting the attentional blink from the no-attentional-blink condition, we identified statistically significantly different sensors and plotted their responses in Fig. 3D. It can be seen that major differences lied in N100 and P300. In either case, the level of response was higher for attentional blink.

**Figure 3.**
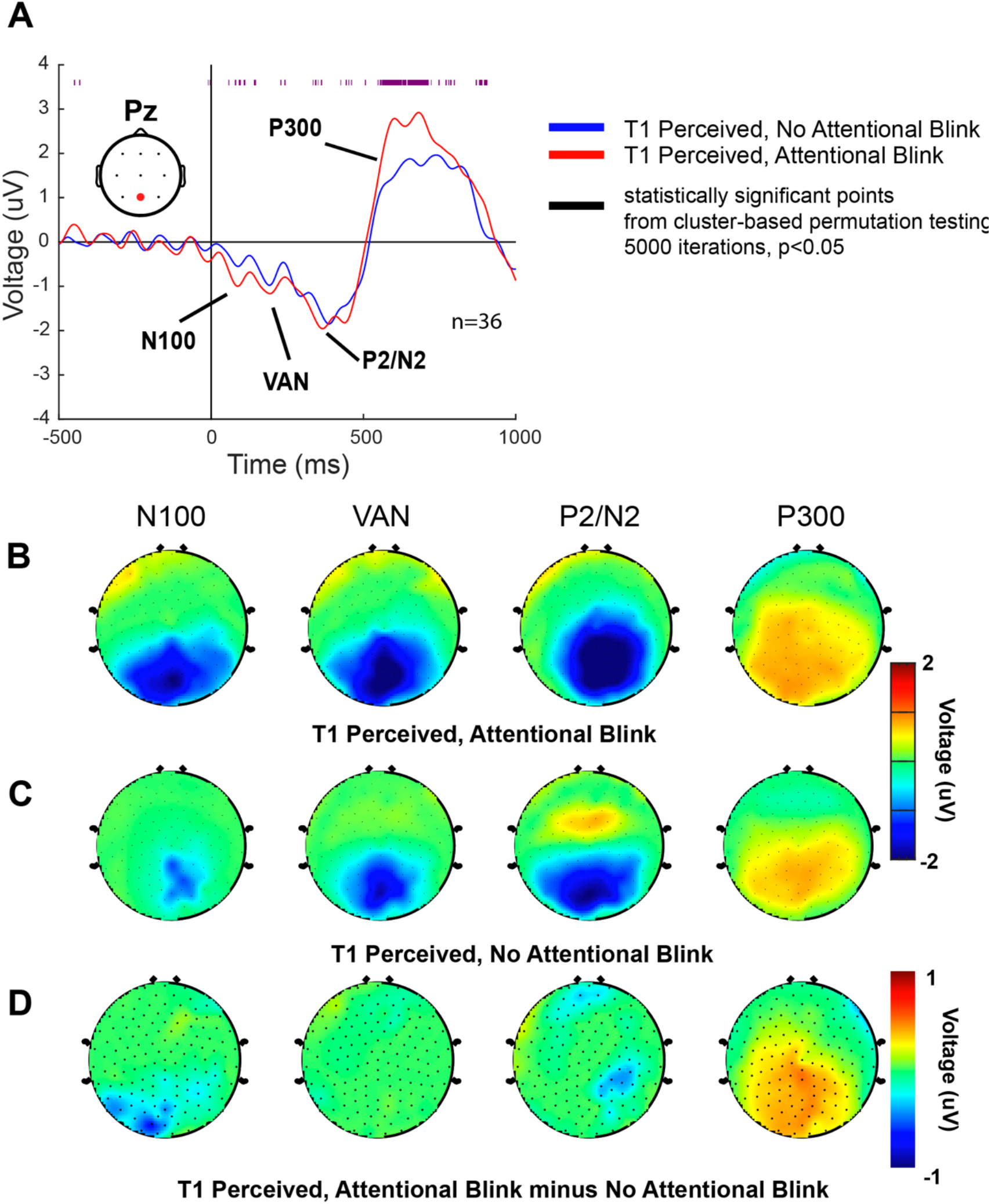
Voltage-space event-related potentials to attentional blink. (A) Comparison of the timecourses of event-related potential between whether attentional blink occurs, given T1 is perceived for pz channel. 200ms and 300ms SOA conditions are combined. Purple bar denotes statistical significance from cluster-based permutation tests (5000 iterations, p<0.05). (B-D)Topolots of voltage responses at 125ms, 250ms, 375ms, and 500ms after T1 onset for the (B) attentional blink occurs, (C) no attentional blink occurs, and (D) attentional-blink minus no-attentional-blink conditions, given T1 is perceived. Only statistically significant sensors from cluster-based permutation tests are included in the topoplots (5000 iterations, p<0.05).

## Methods

### Participants

A total of 62 healthy, adult participants were recruited to participate in this study. Inclusion criteria were: (1) normal with or without correction and (2) normal hearing without an assistive hearing device. Exclusion criteria include: (1) past or current diagnosis of a psychiatric or neurological disorder; (2) vision correction that required glasses.

There were two paradigms: *behavior only* and *behavior+EEG*. 25 subjects participated in the *behavior only* paradigm and 38 subjects participated in the *behavior+EEG* paradigm. 2 subjects in the *behavior+EEG* paradigm were excluded due to corruption of data files.

### Software and Equipment

The behavioral paradigm was developed using Python (www.python.org) and run in the open-source PsychoPy2 environment (https://www.psychopy.org/).

For the *behavior only* group, they viewed task on an MSI Model MS-16H2 laptop running Windows 10 with a 15.6 inch display (screen resolution 1280x780 pixels) and a NVIDIA GeForce graphics card. The experimental laptop was placed on a table at eye-level and centered 85cm from the participant.

For the *behavior+EEG* group, the participants viewed the task on a 17-inch external LCD monitor (Iiyama ProLite E1780SD) mounted on a chart-attached arm mount. The external monitor was positioned at eye-level and centered 55cm from the center of the screen to the participants’ nose bridge.

For both groups, behavioral responses were recorded with a 1x4 inline button response box connected to the experimental laptop via USB and sampled at 1000Hz (Current Designs, Inc.; Model OTR-1x4-L). Regardless of handedness, participants were instructed to make responses with the button response box using their right hand and with fingers sequentially placed along the four buttons, with the first button pressed with the index finger and the fourth button pressed with the pinky finger.

For the *behavior+EEG* group, eye tracking and pupillometry data were collected with the EyeLink 1000 Plus System and software (version 5.09; SR Research, Inc.) running on a Dell PC desktop (Model D13M; Dell, Inc.). The sampling rate was 1000Hz with a 35mm camera lens and infrared illuminator mounted below the task LCD display. EEG data were collected with 257 Ag/AgCl electrodes embedded into an elastic net (Hydrocel GSN 256, Magstim Electrical Geodesics, Inc.) and recorded on a desktop computer (Power Mac G5 Quad; Mac OS X v10.5.8, Apple, Inc.) running NetStation version 4.2.2 (Magstim Electrical Geodesics, Inc. The EEG signal was sampling rate was 1000Hz, amplified with two 128-channel amplifiers high and low-pass hardware filtered at 0.1 and 400Hz, respectively. Signals were acquired as Cz-referenced.

Data analysis was conducted in MATLAB 2020a on Linux (Ubuntu) and MATLAB 2022a on Yale Center for Research Computing’s High Performance Cluster.

### Data Preprocessing

#### EEG data

An automatic pipeline for EEG preprocessing was implemented using the open-source EEG processing toolbox EEGLAB (cite). Data were first 1Hz high-pass filtered. Next, line noise of 60 Hz and 120Hz harmonics were removed. Noisy channels were found and rejected. These channels were restored with spherical interpolation. The data was re-referenced to the common average reference. Epochs of 4001ms duration (2000ms before and 2000ms after each T1 stimulus) were cropped for analysis from the preprocessed session data. The extracted epochs were then concatenated for a 10-component principal component analysis (PCA) to reduce the dimension for ICA training. Then, an independent component analysis (ICA) applied on the PCA decomposed data. ICA components that may be dominated by blinks, eye movements, or heartbeat were automatically detected and removed (cite). Finally, epochs that contained more than 25% bad samples between 200ms pre-T1 and 500ms post-T1 were rejected.

#### Eye-tracking data

Eye-tracking data preprocessing was described in Kronemer et al., 2022[3]. Briefly, blinks and other artifacts were identified using *Stublinks[7]*. Blinks and other artifacts were then removed from the raw pupil timecourses. Interpolation was then used to reconstruct the pupil timecourses. Blink probability was computed as the probability that a blink occurred at each time point across all trials for each subject.

### Statistical Analysis

#### Behavioral Data

Two-sample t-tests were used to compare the differences of T2 detection between T1-percieved and T1-not-perceived conditions. Significance threshold was set at p=0.05.

#### Eye-tracking data

Cluster-based permutation testing was used to assess statistical significance between conditions while controlling for the multiple-comparison problem. The approach and the rationale are detailed in Kronemer et al., 2022.[3] Briefly, a null distribution was generated using 5000 permutations. At each iteration, a two-tailed t-tests were performed independently at each time point. The resulting t-values were clustered based temporal adjacency: i.e., a cluster would be formed if two or more sequential time points were statistically significant. For each cluster, the absolute value of the t-values were summed. The cluster with the largest t-value was retained for that iteration and added to the null distribution. Finally, this procedure was repeated for the original timecourse without permutation. And all resulting cluster that exceed top 5% of the null distribution were considered statistically significant.

#### EEG data

Cluster-based permutation tests as detailed above were used to assess statistical significance. The only difference was that, besides temporal adjacency, spatial adjacency was also used in the clustering step. Specifically, spatial adjacency is defined as all neighboring electrodes for each electrode on the scalp surface. For topopolot visualization, only electrodes that reached statistical significance in cluster-based permutation tests were plotted, while the rest were set to 0.

