## Supplemental Figure 1 for "The Neural Basis of Attentional Blink as a Selective Control Mechanism in Conscious Perception"

**Supp. Info**

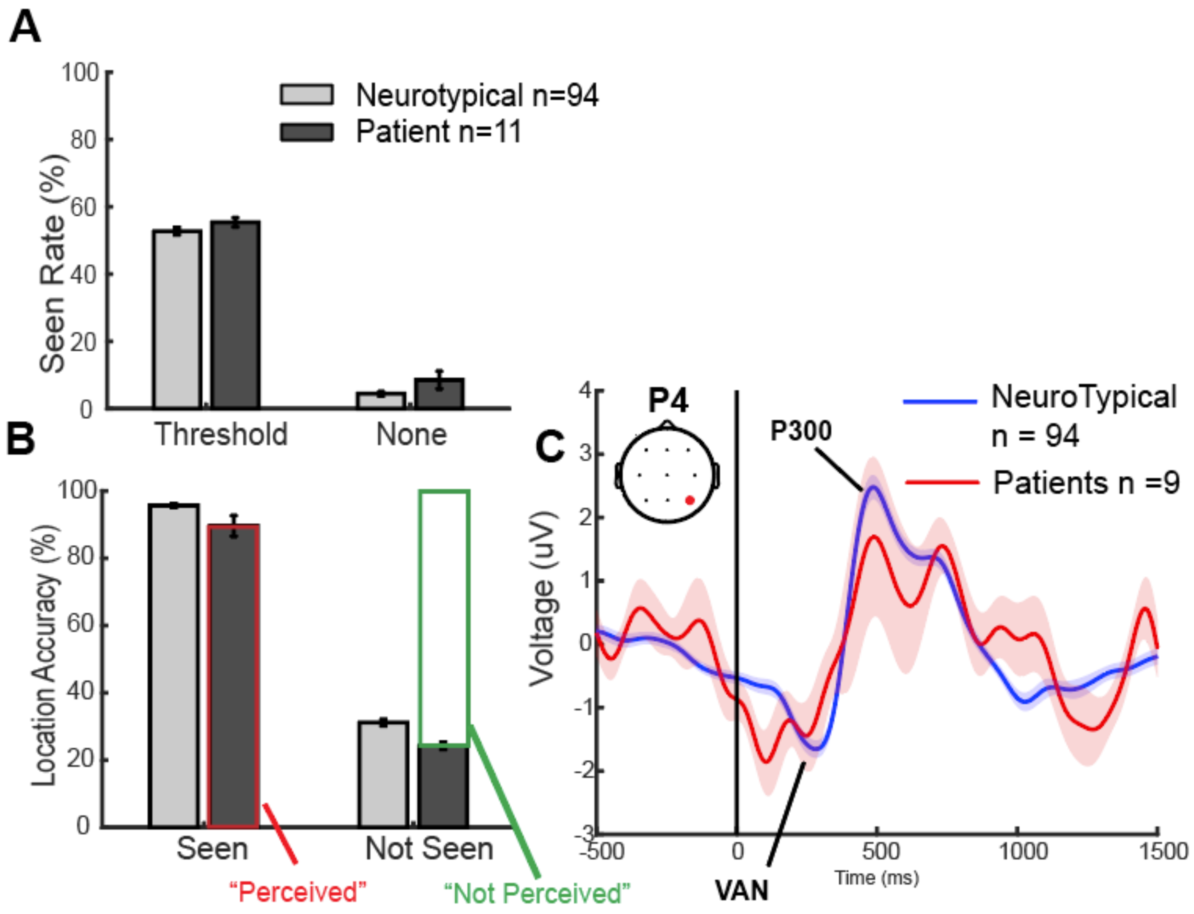


**Supplemental Figure 1.** Neurotypical v. Patients. (A) Percentage of trials where participants report seeing the face when there is a face present at visual threshold compared to blanks (87.5% of stimuli are at-threshold and 12.5% are blanks). (B) Location accuracy when the participants answered they saw or did not see the face, with perception validated by correct location choice. (C) Scalp EEG response (p4 channel) time-locked to face onset (0ms).
